# Second-order regulation: IFN-γ suppresses IL-17A-mediated neutrophilic inflammation

**DOI:** 10.64898/2026.01.05.697792

**Authors:** Vijay Raaj Ravi, Sophia H. Maxfield, Emma N. Niszczak, Hannah Y. Kim, Olivia S. Harlow, Kalyn D. Whitehead, Anukul T. Shenoy

## Abstract

**BACKGROUND:** T helper 1 (T_H_1) cells often accompany T_H_17 cells across diverse tissues in health and disease, including the lungs. However, roles for the T_H_1 effector cytokine, IFN-γ, in T_H_17-driven type 3 inflammation is unclear.

**METHODS:** We devised a reductionistic model to determine the role of IFN-γ in IL-17A-driven inflammation during *Streptococcus pneumoniae* (*Spn*) infection *in vivo.* Briefly, intratracheal instillation of *Spn* along with recombinant TNF-α and IL-17A was used to mimic rapid *Spn*-specific, T_H_17-driven, type 3 inflammation seen in lungs on memory recall infection with *Spn*. Co-instillation of recombinant IFN-γ was used to probe the role for this T_H_1 cell-derived effector cytokine in anti-*Spn* immune response. Immune cellularity in bronchoalveolar lavage (BAL) was used to determine impacts of IFN-γ on type 3 inflammation in murine airways. Mice sufficient for- or lacking-IFN-γ or STAT1 were used to assess the immunoregulatory functions of IFN-γ *in vivo*.

**RESULTS:** IFN-γ promptly muted IL-17A-induced inflammatory cell accumulation in *Spn*-infected airways through a STAT1-dependent mechanism. Both female and male mice demonstrated similar anti-inflammatory effects of IFN-γ on type 3 inflammation. Notably, the impact of IFN-γ was more striking at lower cytokine concentrations. The immunoregulatory effect of IFN-γ against T_H_17-driven type 3 inflammation was also evident in physiologically relevant settings: while immunized wildtype (WT) mice controlled lethal *Spn* infection, immunized IFN-γ knockout mice exhibited even better *Spn* clearance. This heightened antimicrobial resistance, however, was accompanied by overt airway neutrophilia suggesting risk for immunopathology.

**CONCLUSIONS:** Our findings identify a distinct immunoregulatory mechanism that operates within non-lymphoid tissues, where IFN-γ limits IL-17A-mediated type 3 inflammation via STAT1. Thus, the frequent accompaniment of T_H_17 cells with T_H_1 cells may represent a conserved mechanism that restrains immunopathological potential of T_H_17-driven neutrophilic inflammation via STAT1 signaling in non-lymphoid tissues.

## INTRODUCTION

T helper-1 (T_H_1) cells often accompany T_H_17 cells in multiple type 3 inflammatory states across body sites in health and disease (1–4). Some prominent examples include multiple sclerosis in the central nervous system (5), colitis in the colon (6), psoriasis of the skin (7), allergies (8, 9) and microbial infections at barrier epithelial sites (10–16). The reasons for this pairing are incompletely understood. While T_H_1 cell-derived IFN-γ and T_H_17 cell-derived IL-17A inhibit polarization of naïve CD4^+^ T cells to T_H_17 (17) and T_H_1 lineages (18) respectively, several studies have demonstrated that T_H_1 and T_H_17 cells instead synergize to exacerbate disease (2, 19–22). Given these conflicting findings, we sought to resolve the role of IFN-γ using a tractable model of IL-17A-induced type 3 antibacterial inflammation within murine lungs.

*Streptococcus pneumoniae* (*Spn*) is a Gram-positive opportunistic pathogen that frequently colonizes the human nasopharynx and is microaspirated into the lungs of healthy humans globally (23, 24). Despite prolonged spells of colonization and inhalation most healthy individuals clear this pathobiont asymptomatically (25–29). Induction of T_H_17- and T_H_1-polarized lung tissue resident memory (T_RM_) T cells in response to frequent *Spn* inhalation is thought to be a key driver of this robust antibacterial immunity (16, 30). While *Spn*-specific T_H_17 T_RM_ cells, via IL-17A, coax lung epithelial cells to rapidly recruit neutrophils and clear inhaled *Spn* (16, 30), roles of T_H_1 T_RM_ cells and its effector cytokine IFN-γ within *Spn*-experienced lungs remain unclear. This is a major knowledge gap since *Spn-*specific CD4^+^ T_RM_ cells are capable of serotype-independent protection (16) and lend themselves as promising targets for broadly protective vaccines against all serotypes of *Spn;* the latter being a World Health Organization (WHO) priority pathogen in need of better antibiotics and vaccines due to serotype replacement and antibiotic resistance (24, 31–33).

Here, by intratracheally delivering recombinant IL-17A and IFN-γ (along with *Spn*) to immunologically mimic antigen-dependent reactivation of T_H_17- and T_H_1-T_RM_ cells, we explore whether the T_H_1 effector cytokine IFN-γ may alter IL-17A-driven type 3 inflammation of murine airways. By directly delivering the T cell-effector cytokines to the lung mucosa, we bypass the first-order cross-regulation between T_H_1 and T_H_17 cells that occurs within secondary lymphoid organs (SLOs) (17, 18) to uncover a second-order cross-regulatory mechanism wherein IFN-γ limits IL-17A-mediated type 3 inflammation in a STAT1-dependent manner within the airways. Thus, our findings suggest that T_H_1 biology may have evolved to accompany T_H_17 cells to limit their immunopathological potential at non-lymphoid tissue (NLT) sites.

## RESULTS

### IFN-γ decreases IL-17A-driven neutrophilia in a sex-independent manner

Recurrent inhalation of *Spn* induces T_H_17- and T_H_1-polarized T_RM_ cells within experienced lungs (16, 30). While T_H_17 T_RM_ cells rapidly recruit neutrophils to drive robust antimicrobial immunity (14, 16, 34), roles of T_H_1 T_RM_ cells within *Spn*-experienced lungs remain unclear. We sought to investigate whether the T_H_1 effector cytokine IFN-γ plays any roles in modulating IL-17A-induced type 3 inflammation within such *Spn*-experienced murine lungs. Informed by knowledge T_H_1 and T_H_17 T_RM_ cells coexist in *Spn*-experienced lungs at comparable frequencies (16, 30) and secrete similar levels of IFN-γ or IL-17A in response to *Spn* reencounter, respectively (16, 30), we developed a reductionist *in vivo* model to mimic rapid anti-*Spn*-memory recall. Briefly, naïve C57BL/6J mice were intratracheally administered a cocktail of 200ng of recombinant TNF-α and IL-17A (along with a non-lethal isolate of *Spn* belonging to serotype 19F, Sp19F) to mimic rapid T_H_17 T_RM_ response against homeostatic inhalation of *Spn* (**Fig 1A**). Immune cellularity of the airways was assessed in bronchoalveolar lavage (BAL) 8 hours post-infection when T_H_17 T_RM_ cell-driven airway neutrophilia is robust within *Spn*-experienced murine lungs (16, 34). Of note, instilled cytokine cocktails included- or lacked-200ng of recombinant IFN-γ to model presence or absence of *Spn*-specific T_H_1 T_RM_ cells respectively. The treatment group that received IFN-γ will hereafter be referred to as “+ IFN-γ” whereas the group without IFN-γ will be called “-IFN-γ”. By delivering the T_H_ effector cell cytokines directly at the lung mucosa, this model bypasses the first-order cross-regulation between T_H_1 and T_H_17 cells that occurs within SLOs (17, 18) and lends a powerful system to probe for roles of IFN-γ in altering IL-17A-driven type 3 inflammation in the lung mucosa where comparable numbers of T_H_1 and T_H_17 T_RM_ cells reside post their egress from SLOs (16, 30). Consistently, both female (**Fig 1B**) and male mice (**Fig 1C**) co-administered IFN-γ showed reduced accumulation of total, polymorphonuclear (PMNs), and mononuclear cells compared to their “-IFN-γ” counterparts (**Fig 1D**, compiled and normalized). Thus, our data suggest that the T_H_1 cytokine IFN-γ reduces IL-17A-driven influx of PMNs and mononuclear cells in mouse airways, independent of sex.

**Figure 1:**
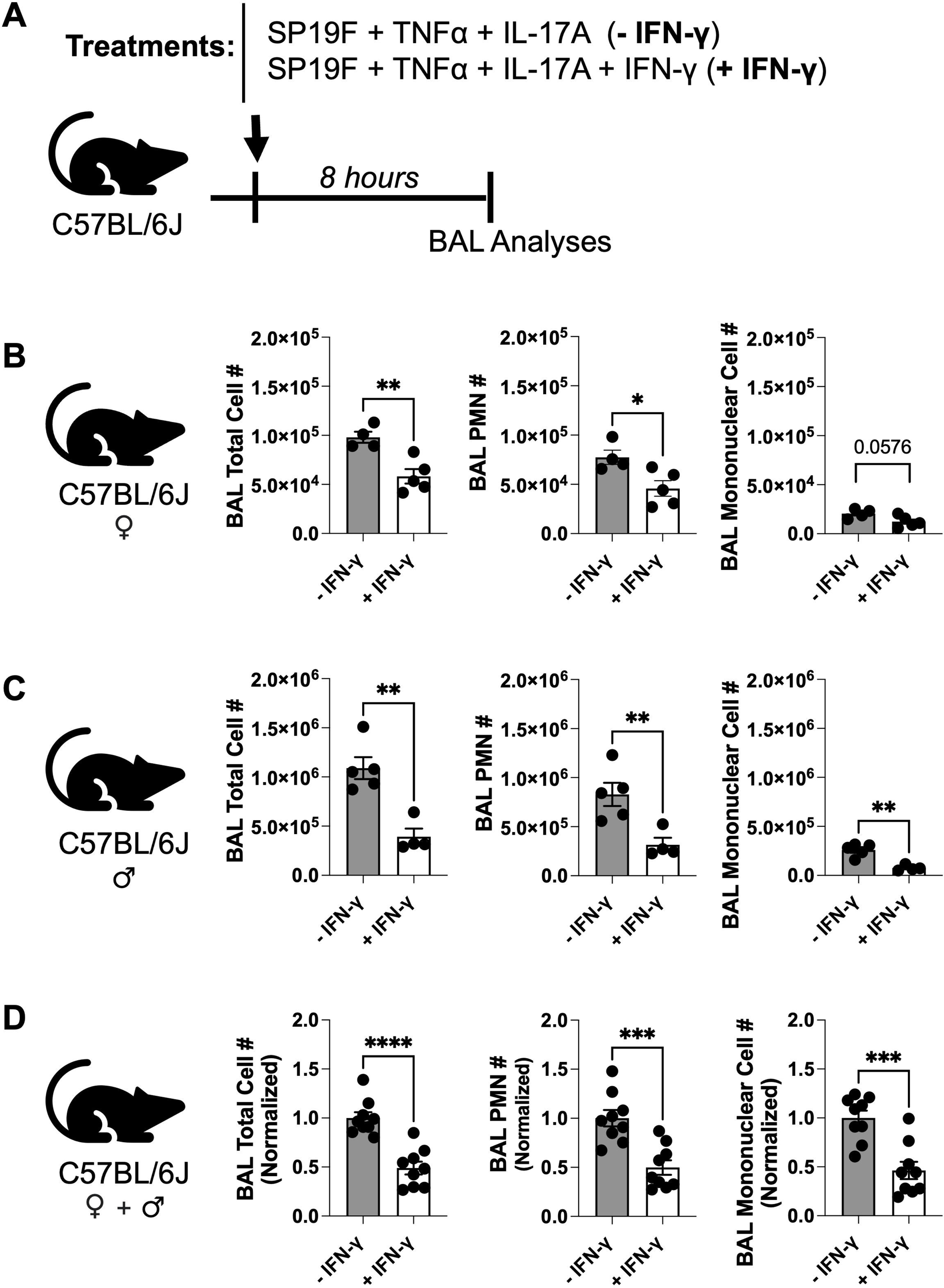
IFN-γ dampens IL-17A-driven airway inflammation in a sex-independent manner. **A)** Schematic of experimental model used. **B-D)** Total cell number, PMN number, and mononuclear cell number present in the bronchoalveolar lavages (BAL) 8 hours post treatment in **(B)** Female wildtype (WT) mice, **(C)** Male WT mice, and **(D)** Aggregated female and male WT mice data normalized to (-IFN-γ) treatment group within each sex. Unpaired t test. *p* value: ns≤ 0.1234, *≤ 0.0332, **≤ 0.0021, ***≤ 0.0002, ****≤ 0.0001. All data have n≥4 mice, 2 experiments, mean ± SEM.

### IFN-γ exerts stronger immunomodulatory effects at lower doses

Multiple cytokines function in a concentration-dependent manner where higher levels of the cytokines exert stronger biological effects (35, 36). Given that the 200ng doses we used are much higher than the physiologically relevant levels of these cytokines that accumulate in *Spn*-experienced lungs in response to memory recall (16), we asked if reducing the levels of these cytokines would extinguish the immunoregulatory effect of IFN-γ. We intratracheally administered cocktails of 50ng, 100ng, or 200ng of each cytokine along with Sp19F followed by analyses for BAL cellularity (**Fig 2A**). Surprisingly, the immunoregulatory effects of IFN-γ were markedly enhanced at lower concentrations (**Fig 2B-G**). While 50ng IFN-γ led to a 5.2-fold decrease in BAL total cellularity, 200ng instead exhibited only a 1.7-fold decrease **(Fig 2B,E)**. A similar dose-dependent effect was observed for both PMN (**Fig 2C,F**) and mononuclear cell accumulation in the airways (**Fig 2D,G**). Thus, IFN-γ becomes a potent modulator of IL-17A-driven airway immune cellularity at lower cytokine concentrations. Given this stark effect, we hereafter use 50 ng cocktails for the rest of the study unless specified otherwise.

**Figure 2:**
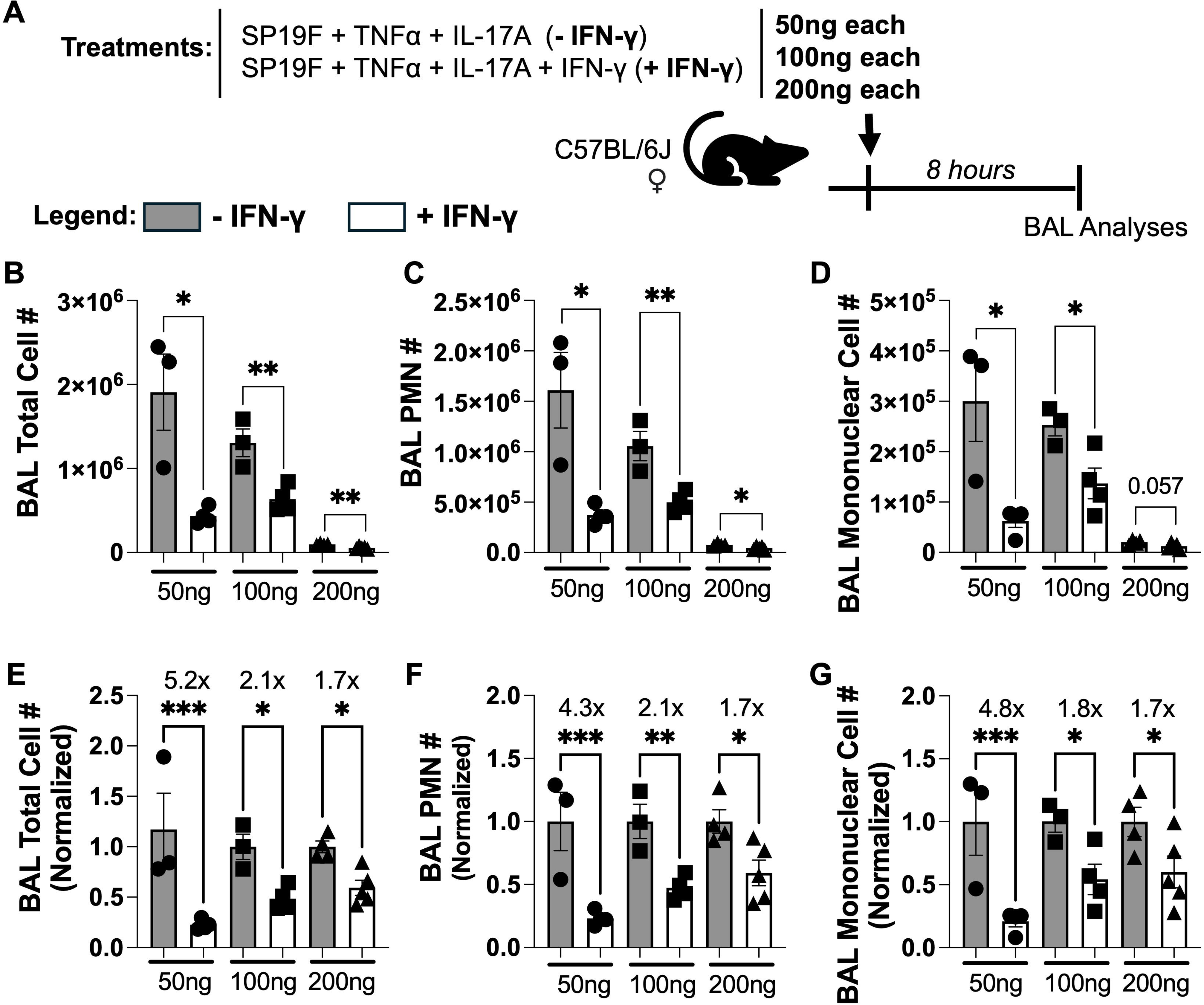
Lower doses of IFN-γ exhibit greater immunoregulatory effects. **A)** Schematic of experimental model used. **B)** Total cell number, **C)** PMN cell number, and **D)** Mononuclear cell number in BALs from female WT mice 8 hours post treatment with varying cytokine concentrations. **E)** Total cell number, **F)** PMN cell number, and **G)** Mononuclear cell number in BALs from female WT mice 8 hours post treatment with varying specified concentrations normalized to (-IFN-γ) treatment group. Unpaired t test. *p* value: ns≤ 0.1234, *≤ 0.0332, **≤ 0.0021, ***≤ 0.0002, ****≤ 0.0001. All data have n≥3 mice, 2 experiments, mean ± SEM.

### IFN-γ broadly mutes type 3 inflammatory cell landscape

IL-17A-driven neutrophilic inflammation is accompanied by enhanced accrual of Ly6C^+^ monocytes (37) and macrophages in the inflamed tissue (38). To assess the extent of immunoregulatory effect of IFN-γ on IL-17A-driven inflammatory cell landscape, we performed flow cytometry on BAL (**Fig 3A**, Gating Strategy in **Supplementary Fig S1**). In addition to reducing total cellularity (**Fig 3B**) and neutrophils (**Fig 3C**), IFN-γ consistently reduced the numbers of multiple type 3-inflammatory cells including tissue resident alveolar macrophages (TRAMs) (**Fig 3C**), Ly6C^+^ monocytes (**Fig 3D**), and monocyte-derived alveolar macrophages (MoAMs) (**Fig 3E**) that accumulated within inflamed airways of mice independent of sex. Beyond the aforementioned cell types, we also expanded our analyses to include other key innate immune cell subsets that may be impacted by IFN-γ. While IFN-γ impacted accumulation of Ly6C^-^ monocytes **(Supplementary Figs S2F and S2M)**, CD11b^+^Ly6C^-^ dendritic cells (CD11b^+^ DCs) **(Supplementary Figs S2D and S2K)** and CD11b^-^Ly6C^+^ DCs **(Supplementary Figs S2B and S2I)** in sex-dependent fashion, it consistently decreased numbers of CD11b^+^Ly6C^+^ DCs **(Supplementary Figs S2C and S2J)** and conventional type 1 DCs (cDC1s) **(Supplementary Figs S2A and S2H)** across both sexes of mice **(Supplementary Figs S2A,B)**. Numbers of eosinophils and lymphocytes remained unaffected. Taken together, our data suggest that IFN-γ substantially remodels the inflammatory cell landscape of murine airways experiencing type 3 inflammation.

**Figure 3:**
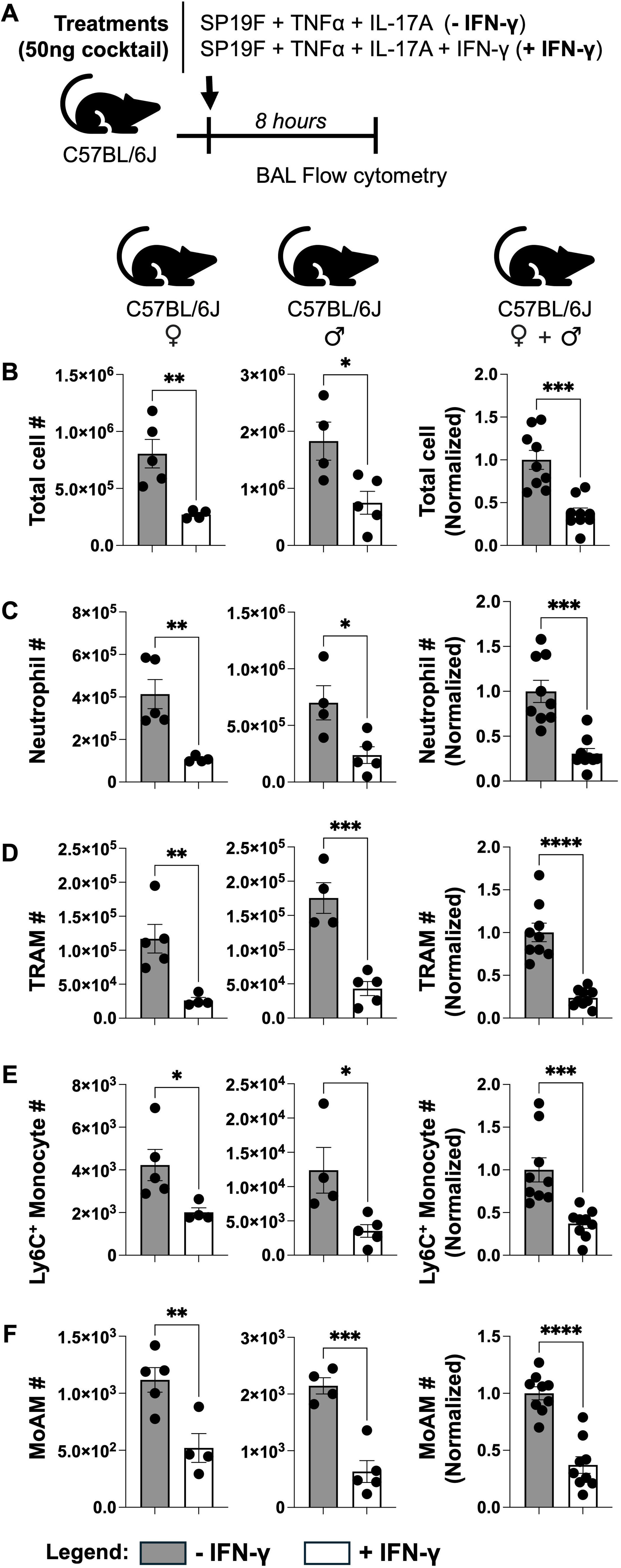
IFN-γ broadly dampens the type 3 inflammatory cell landscape. **A)** Schematic of experimental model used. **B)** Total cellularity, **C)** Neutrophils, **D)** TRAMs, **E)** Ly6C+ Monocytes, and **F)** MoAMs in BALs of female and male WT mice 8 hours post treatment. Fold changes in aggregated data were calculated relative to the -IFN-γ treatment group within each sex. Unpaired t test. *p* value: ns≤ 0.1234, *≤ 0.0332, **≤ 0.0021, ***≤ 0.0002, ****≤ 0.0001. All data have n≥3 mice, 2 experiments, mean ± SEM.

### IFN-γ signaling via STAT1 inhibits type 3 inflammation

IFN-γ conventionally signals via the Janus kinase (JAK)–signal transducer and activator of transcription 1 (STAT1) pathway to exert its effects (39). We sought to determine whether STAT1 is required for IFN-γ’s immunoregulatory activity within lung mucosa during type 3 inflammation (**Fig 4A**). To determine this, we intratracheally instilled cocktails of all three cytokines at 50 ng each (along with Sp19F) into wildtype (WT) and STAT1 knockout mice and measured airway cellularity using flowcytometry on BAL. Consistently, all the key inflammatory cell types associated with type 3 airway inflammation were elevated in airways of both female and male STAT1 KO mice despite IFN-γ instillation **(Fig 4B–4F)**. In addition to these, numbers of CD11b^+^Ly6C^+^ DCs **(Supplementary Figs S3D and S3K)** and cDC1s **(Supplementary Figs S3A and S3H)** were also elevated in mice lacking STAT1 independent of their sex **(Supplementary Figs S3A,B)**. This is consistent with their observed dependence on IFN-γ during type 3 inflammation **(Supplementary Figs S2A,B)**. Thus, our data suggests that STAT1 deficiency phenocopies overt type 3 inflammation displayed by WT mice lacking IFN-γ instillation. Indeed, post-hoc concatenation of data from WT mice receiving 50ng of TNF-α and IL-17A (i.e. - IFN-γ WT) with data from WT and STAT1 KO mice receiving 50ng of all three cytokines (i.e. +IFN-γ WT and +IFN-γ STAT1 KO), followed by Principal Component Analysis (PCA) to reduce the dimensionality of our data revealed that STAT1 KO mice clustered close to the “-IFN-γ WT” group and away from “+IFN-γ WT” group **(Figure 4G)**. We next queried the cell types that influenced the direction and magnitude of each principal component. Among all the cell types identified, PCA loading plot revealed that the major type 3 inflammatory cells (namely neutrophils, TRAMs, Ly6C^+^ monocytes, and MoAMs) drove the co-clustering towards the top left quadrant **(Figure 4H)** reflecting strong positive correlations and indicating their primary contribution in driving separation of STAT1 KO mice towards “-IFN-γ WT” and away from “+IFN-γ WT” mice in the PCA. Other contributors to the clustering appeared to be eosinophils and cDC1s, suggesting their dependence on IFN-γ-STAT1 signaling during type 3 inflammation (**Fig 4H, S2, S3**). Collectively, these findings demonstrate that IFN-γ mediates its immunoregulatory effects on lung mucosa during type 3 inflammation in a STAT1-dependent manner.

**Figure 4:**
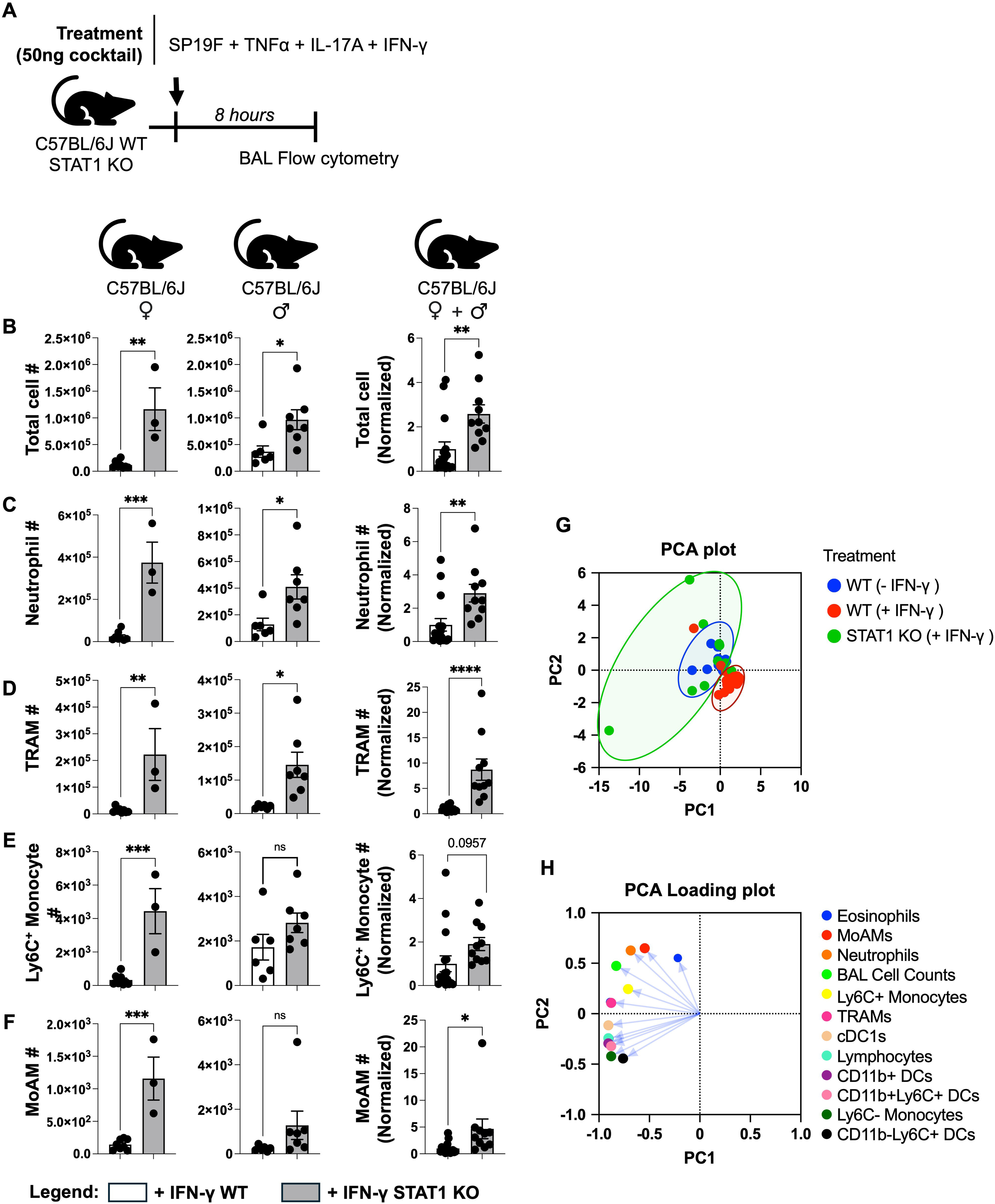
IFN-γ suppresses type 3 inflammation through STAT1. **A)** Schematic of experimental model used. **B)** Total cell number, **C)** Neutrophil, **D)** TRAMs **E)** Ly6C+ Monocytes, and **F)** MoAMs in BALs of female and male WT and STAT1 KO mice 8 hours post treatment. Fold changes in aggregated data were calculated relative to the -IFN-γ treatment group within each sex. Unpaired t test. **G)** Principal component analysis (PCA) plot of compiled BAL cellularity data from C57BL/6J WT and STAT1 KO mice based on total cell numbers and 11 distinct cell types identified by flow cytometry. **H)** PCA loadings indicating the magnitude and direction of each variable’s contribution to a principal component. *p* value: ns≤ 0.1234, *≤ 0.0332, **≤ 0.0021, ***≤ 0.0002, ****≤ 0.0001. All data have n≥3 mice, 2 experiments, mean ± SEM.

### IFN-γ impairs bacterial clearance while restricting neutrophilic inflammation

Lung-resident T_H_17 T_RM_ cells use IL-17A to coax local epithelial cells (34) and fibroblasts (40) to rapidly recruit neutrophils within 7-8 hours of memory recall to confer robust antimicrobial immunity in the lungs (16, 34). We asked if T_H_1 T_RM_ cell-derived IFN-γ may dampen this T_H_17-driven protective inflammation. We intratracheally administered Sp19F into WT and IFN-γ-deficient (IFN-γ KO) mice on days 0 and 7 to generate T_RM_ cells within the lungs (16, 34); control WT mice received sterile PBS. Following 4 weeks of rest, all groups were intratracheally challenged with a lethal serotype mismatched strain of *Spn* belonging to serotype 3 (Sp3) and bacterial clearance was assessed (**Fig 5A**). Strikingly, while experienced WT mice cleared Sp3 by 100-fold within 24 hours of infection, the experienced IFN-γ KO mice cleared the bacteria by ∼5000-fold within 24 hours **(Fig 5B)**. This enhanced microbial clearance was accompanied by greater neutrophil accumulation within airways of *Spn-*experienced IFN-γ KO mice compared to *Spn-*experienced IFN-γ sufficient WT mice **(Fig 5C)**. Taken together, our data suggest that IFN-γ secreted by T_H_1 T_RM_ cells may restrict T_H_17 T_RM_ cell-driven neutrophilic inflammation to compromise robust antimicrobial immunity in experienced lungs.

**Figure 5:**
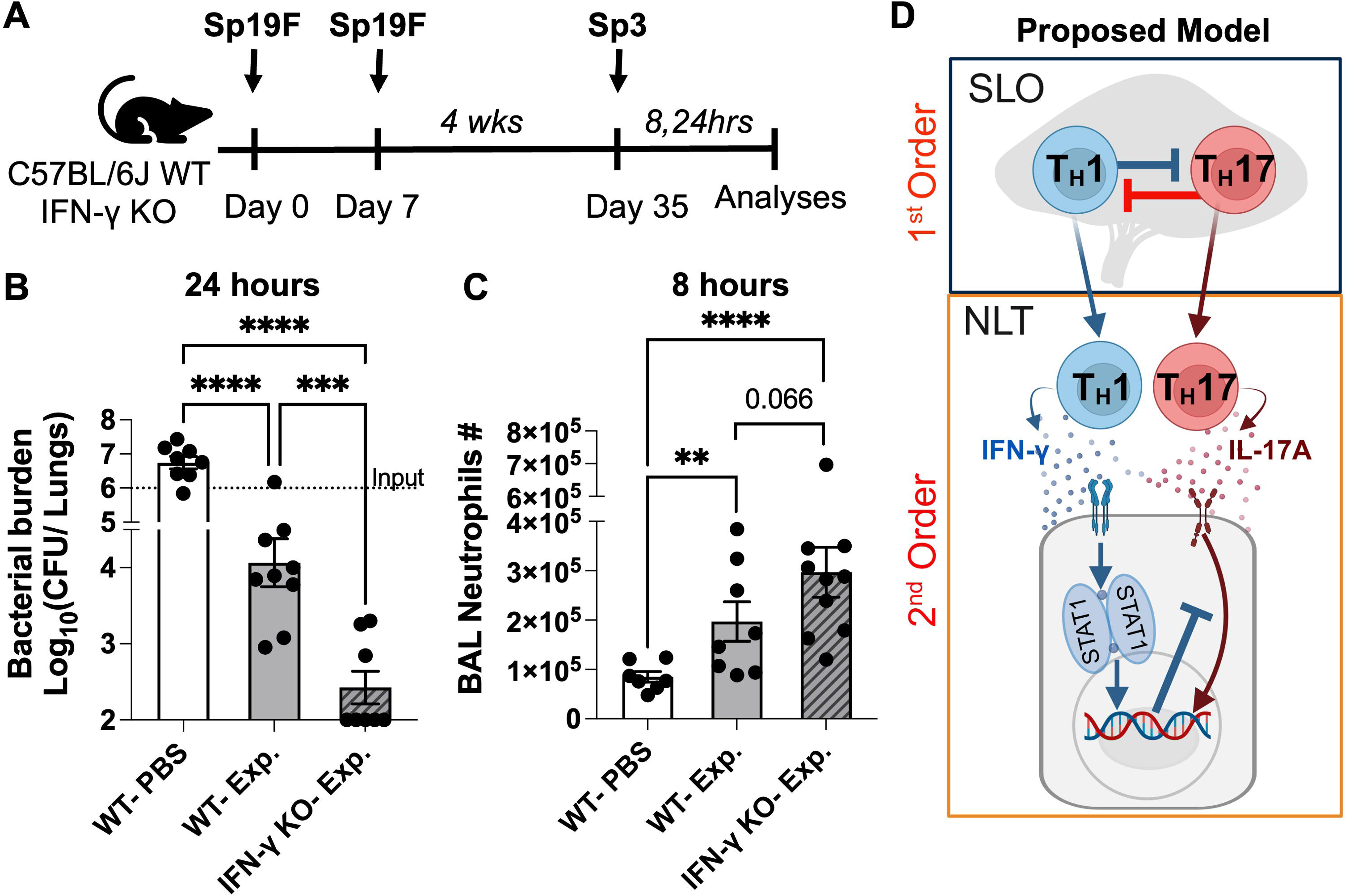
IFN-γ limits neutrophilic inflammation and compromises bacterial clearance on memory recall. **A)** Schematic of experimental model used. **B)** *Spn* serotype 3 CFU counts 24 hours post infection (hpi). Dotted line indicates the bacterial load in the challenge inoculum. Ordinary one-way ANOVA. **C)** Neutrophil numbers in BAL of naive and *Spn* experienced WT and IFN-γ KO mice 8 hpi. Lognormal ordinary one-way ANOVA. **D)** Graphical abstract illustrating our proposed working model of second-order cross-regulation between type 1 and type 3 inflammatory cytokines within non-lymphoid tissues. *p* value: ns≤ 0.1234, *≤ 0.0332, **≤ 0.0021, ***≤ 0.0002, ****≤ 0.0001. All data have n≥3 mice, 2 experiments, mean ± SEM.

## DISCUSSION

Herein, we directly delivered recombinant IFN-γ and IL-17A cytokines at the lung mucosa to find that the T_H_1 effector cytokine IFN-γ limits IL-17A-mediated type 3 inflammation in a STAT1-dependent- and sex-independent-manner; these regulatory effects of IFN-γ were more pronounced at lower cytokine concentrations. We find that IFN-γ potently inhibited accumulation of various myeloid cells involved in type 3 inflammation including tissue resident alveolar macrophages (TRAMs), neutrophils, Ly6C⁺ monocytes, monocyte-derived AMs (MoAMs), cDC1s, and CD11b^+^Ly6C^+^ DCs. Finally, using *Spn*-experienced mice that lack IFN-γ, we demonstrate that IFN-γ restricts T_H_17 T_RM_ cell-driven neutrophilic inflammation to dampen antibacterial immunity in experienced lungs.

T_H_1 cells consistently accompany T_H_17 cells across multiple body sites during various states of health and disease (1–4) ranging from infections (10–16, 41–43), autoimmunity (1, 44, 45), allergy (8, 9, 46, 47), and fibrosis (48, 49) in both mice and humans. However, the significance of this pairing beyond SLOs where they inhibit polarization of naïve T cells into the opposite lineage (17, 18) has remained unclear. Our data suggests that T_H_1 cells, via their key effector cytokine IFN-γ, regulate T_H_17-driven type 3 inflammation within the non-lymphoid tissues. These findings partially explain why mice deficient in type 1 immunity consistently have worse outcomes in myriad models of type 3 inflammatory disease including experimental autoimmune encephalitis (EAE) model of MS (50), tuberculosis (51, 52), bacterial pneumonia (53) and allergies (8, 54). T_H_17 cells incite their inflammatory activities by stimulating local epithelial cells (34) and fibroblasts (40) to produce neutrophil attracting chemokines via IL-17 signaling. Given that mere instillation of recombinant IFN-γ directly into naive lungs was sufficient to rapidly mute IL-17A-driven inflammation, our data suggest that IFN-γ may exert its immunoregulatory effect via an extra-lymphocytic mechanism that involves signaling into tissue resident cells within non-lymphoid tissues. We propose that pairing of T_H_17 cells with T_H_1 cells represents an evolutionarily conserved strategy to ensure just enough type 3 inflammation to control the antigenic insult while simultaneously imposing restraints on the immunopathological potential of T_H_17 cells. The price that our immune system pays for this measured type 3 inflammation is more modest antimicrobial clearance which can only reach its full potential when IFN-γ-mediated constraints are lifted. Supporting this, previous studies with *Mycobacterium tuberculosis* show that mice lacking IFN-γ receptor experienced T_H_17-driven excessive neutrophilic infiltration and lethal immunopathology within infected lungs (52); mere depletion of neutrophils in these mice was sufficient to improve survival suggesting that the overt immunopathology, and not uncontrolled bacteria, was the cause of death. Thus, efforts towards developing T_H_17 T_RM_ cell-directed vaccines should consider the yin-yang relationship between type 1 and type 3 cytokines and should consider promoting T_H_1 T_RM_ cells concomitantly to balance antimicrobial immunity and risk for immunopathology. Taken together, our findings fundamentally extend our knowledge regarding cross-regulation between type 1 and type 3 inflammatory cells beyond the SLOs to identify a distinct, second-order regulatory mechanism that operates within non-lymphoid tissues where tissue resident cells integrate type 1 and type 3 effector cytokine signals to fine-tune inflammation **(Figure 5D)**. We anticipate this tissue resident cell to be a non-myeloid cell since a recent study found that myeloid cell STAT1 was dispensable for regulation of type 3 neutrophilic inflammation in a *Klebsiella pneumoniae* driven bacterial pneumonia (53). Precisely how IFN-γ signaling mutes IL-17A-driven inflammation and in which tissue resident cell/s is an active area of investigation with significant translational and clinical potential.

Our results find that IFN-γ signaling suppresses type 3 inflammation via its conventional transcriptional factor STAT1. The significance of STAT1 in suppressing IL-17-driven inflammation is evident from clinical observations that individuals with STAT1 gain-of-function (GOF) mutations are unable to control mucosal infections by common pathobionts such as *Candida albicans* (55, 56)*, S. pneumoniae* (57, 58)*, Staphylococcus aureus* (57, 58)*, Pseudomonas aeruginosa* (57, 58)*, and Haemophilus influenzae* (57, 58); all agents that require T_H_17 derived IL-17 and resulting type 3 inflammation for timely control. While the role for lymphocyte-intrinsic STAT1 in precluding protective T_H_17 cell development within SLOs of these patients is well recognized (17), our findings suggest an additional extra-lymphocytic layer of dysregulation where overt STAT1 activation within tissue resident cells of such patients may mute neutrophil recruitment to render type 3 inflammation insufficient for microbial control at mucosal barriers. Indeed, >25% of patients with STAT1 GOF function mutations who receive hematopoietic stem cell transplantation (HSCT, to normalize immune-cell intrinsic STAT1) die from infections within few years post-HSCT (58). Instead, broad inhibition of IFN signaling with JAK inhibitors improve symptoms and lead to complete recovery in such patients (58, 59). It is tempting to posit that JAK inhibitors outperform HSCT owing to their ability to agnostically inhibit IFN signaling within hematopoietic and non-hematopoietic cells; the latter would be unaltered by HSCTs.

IFN-γ displays enhanced immunomodulatory effects at lower cytokine concentrations. Previous studies have found that lower concentrations of IFN-γ compromises the induction of optimal antimicrobial activity of neutrophils (35), as well as natural killer cells and T cells (36). Our data instead find that raising the concentration of cytokines weakened the inhibitory effect of IFN-γ on type 3 inflammatory cell infiltrates from 5.2-fold (for 50ng) to 1.7-fold (for 200ng) per mouse. Our findings thus suggest that IFN-γ’s immunoregulatory activity may be at its most potent at lower concentrations that are closer to physiological levels produced during healthy immunity which is in stark contrast to IFN-γ’s immunostimulatory activity that may require high concentrations of the cytokine (35, 36). This also implies that excessive IFN-γ (such as those seen in severe or chronic diseases) may possess little to no ability to mute T_H_17-driven type 3 inflammation but may instead synergize with IL-17A to worsen immunopathology by compromising epithelial barrier integrity (60, 61) and overtly activating recruited innate and adaptive immune cells in diseases such as inflammatory bowel disease (19, 62), autoimmune thyroiditis (63, 64), diabetes (65), among others. Thus, in addition to highlighting the importance of hormesis as a key determinant of IFN-γ-driven biology, our findings encourage caution with use of- and interpretation of data from-experiments that employ high levels of recombinant IFN-γ to study impacts of IFN-γ signaling may reflect chronic pathological conditions. Direct comparative studies assessing effects of IFN-γ at concentrations lower than 50ng and higher than 200ng in our model are now warranted.

Beyond IFN-γ, our studies have implications for type 1 (IFN-I) and type 3 IFN (IFN-III) biology as well, owing to their dependence on STAT1 (66, 67). While their STAT1-dependent inhibitory effects at low doses may explain IFN-I and IFN-III’s suppressive effects of neutrophilic inflammation in some contexts (68–70), their weakened immunoregulatory activity at high doses may explain their starkly contrasting and detrimental effects during severe/chronic infections (71–73) and autoimmunity (74). It is tempting to posit that patients with autoantibodies against IFN-I (and thus insufficient STAT-1 signaling) experience worsened SARS-CoV-2 driven airway neutrophilia leading to overt lung immunopathology and severe disease (75, 76).

Taken together, using a reductionist model of lung-resident *Spn*-specific T_RM_ cell memory response that bypasses first-order T_H_1 and T_H_17 cell cross-regulation in SLOs (17, 18), we uncover a second-order, cross-regulatory mechanism where IFN-γ limits IL-17A-mediated type 3 inflammation via STAT1 within non-lymphoid tissues. Our findings thus suggest that T_H_1 biology may have evolved to accompany T_H_17 cells to limit their immunopathological potential at peripheral tissue sites. This expanded view of cytokine cross-regulation highlights the importance of context, site, and dose in immune-modulation, and opens new avenues for next generation therapeutic strategies that will finely balance protective and pathological inflammation at barrier tissue sites.

## MATERIAL AND METHODS

### LEAD CONTACT AND MATERIALS AVAILABILITY

Correspondence and requests for information, resources and reagents should be directed to Anukul T. Shenoy. All data will be made available by the corresponding authors upon reasonable request.

### Mice

6-week-old C57BL/6J (stock# 000664), IFN-γ KO (B6.129S7-Ifng^tm1Ts^/J, Stock# 002287), and STAT1 KO (B6.129S(Cg)-Stat1^tm1Dlv^/J, Stock# 012606) mice were obtained from Jackson labs (USA) and subsequently bred in-house to ensure acclimation to local conditions. Mice aged 7–14 weeks were used for all experiments. Animals were housed in a specific pathogen-free facility on a 12-hour light/dark cycle with ad libitum access to standard chow and water. Euthanasia was performed using isoflurane overdose, and death was confirmed via pneumothorax prior to organ collection. All animal procedures conformed to the guidelines approved by the Institutional Animal Care and Use Committee at the University of Michigan, Ann Arbor.

### Intratracheal cytokine and *Streptococcus pneumoniae* administration

Mice were administered 200ng, 100ng, or 50ng of recombinant mouse TNF-α (315-01A-20UG, Peprotech), IL-17A (Cat# 210-17-25UG, Peprotech), and/or IFN-γ (Cat# 315-05-100UG, Peprotech) along with ∼10^6^ CFU of serotype 19F *Spn* (Sp19F, Strain EF3030) in 100μL PBS for intratracheal instillation models mimicking antigen-experienced mice.

### Mice CFU experiments

*Spn* experienced mice were generated as previously described (16, 34). Mice were anesthetized via isoflurane and were intratracheally infected with either 1-5 × 10^6^ CFU of serotype 19F *Spn* (Strain EF3030) suspended in 100μL of sterile PBS or just 100μL of sterile PBS (control/naïve group) on days 0 and 7 and then allowed to recover for 35 days. On day 36, the mice were intratracheally challenged with 1 × 10^6^ CFU of serotype 3 *Spn* (Sp3, ATCC 6303), suspended in 100μL of sterile PBS. 24 hours after the challenge the mice were euthanized using isoflurane overdose and the lungs were used to enumerate CFUs.

### Bronchoalveolar lavage (BAL) collection and analyses

Euthanized mice were exsanguinated, and their tracheas cannulated with an 18-gauge cannula followed by 8 rounds of lavage with sterile PBS. The cell pellets from BALs were used to perform cytospins or flowcytometry, and airway cellularity enumeration.

### Bronchoalveolar lavage cytology

Cell pellets were deposited on Fisherbrand™ Superfrost™ Plus Microscope Slides (Cat# 12-550-15, ThermoFisher Scientific) for cytospins using a cytocentrifuge. Slides were air-dried overnight before staining with Hema 3™ Stat Pack solutions. The staining protocol involved dipping the slides for 6 seconds in fixative, 12 seconds in solution I, and 12 seconds in solution II, followed by rinsing with deionized water. Slides were then air-dried for an additional day prior to imaging with an Olympus BX60 microscope.

### Flow cytometry

Flow cytometric analysis of diverse cell types in the BAL from mice was performed using the BD LSRFortessa™ Cell Analyzer. The antibodies used to stain the samples are i) APC anti-mouse/human CD11b (Clone M1/70) - Biolegend Cat# 101212, RRID:AB_312795; ii) PE-Cy7 anti-mouse CD11c (Clone N418) – Biolegend Cat# 117318, RRID:AB_493568; iii) Alexa Flour 700 anti-mouse CD45 (Clone 30-F11) – Biolegend Cat# 103128, RRID:AB_493715; iv) PE anti-mouse Ly-6G (Clone 1A8) – Biolegend Cat# 127608, RRID:AB_1186099; v) Ly-6C Monoclonal Antibody (Clone HK1.4) – Invitrogen Cat# 48-5932-82, RRID:AB_10805519; vi) APC-Cy™7 Rat Anti-Mouse Siglec-F (Clone E50 – 2440) – BD Pharmingen Cat# 565527, RRID:AB_2732831, vii) 7-AAD – BD Pharmingen Cat# 559925, RRID:AB_2869266. The data was analyzed with FlowJo software (BD Biosciences). Gating strategies are detailed in the Supplementary Figures and were established using Fluorescence-Minus-One (FMO) controls.

### Statistical Analyses

Statistical analyses were conducted using Prism 10 (version 10.3.0, GraphPad). Statistical significance was defined as a p value depicted in GraphPad style. Details regarding sample size, number of experimental replicates, and statistical tests applied are provided in each figure legend. Data are presented as mean ± SEM for all figures, except for supplementary figure **S1**, which display flow cytometry gating strategy.

## ACKNOWLEDGEMENTS

We thank the Dept. of Microbiology and Immunology at the University of Michigan for support with flow cytometry. We acknowledge thank Drs. Bethany Moore, and Denise Kirschner for their valuable comments on the manuscript. This work was supported by NIH grants T32AI007413 (supports S.H.M. and K.D.W.), T32AI007528 (supports O.S.H.), T32TR004371 (supports H.Y.K.), R00HL157555 to A.T.S. as well as by the Institutional funding and Endowment for Basic Science from University of Michigan to A.T.S.

## AUTHOR CONTRIBUTIONS

Conceptualization, A.T.S.; Experimental Design and Interpretation, V.R.R. and A.T.S.; Investigation, V.R.R., S.H.M, E.N.N, H.Y.K., O.S.H., K.D.W., and A.T.S.; Validation, V.R.R. and A.T.S.; Supervision, A.T.S.; Project Management, V.R.R. and A.T.S.; Software, V.R.R. and A.T.S.; Formal Analysis, V.R.R. and A.T.S.; Data Curation, V.R.R. and A.T.S.; Visualization, V.R.R. and A.T.S.; Resources V.R.R. and A.T.S.; Writing-Original Draft, V.R.R.; Writing-Review & Editing, V.R.R., S.H.M, E.N.N, H.Y.K., O.S.H., K.D.W., and A.T.S.

## SUPPLEMENTAL FIGURES AND FIGURE LEGENDS

**Figure S1:** Flow cytometry gating strategy for distinguishing various myeloid cell populations

**Figure S2: Extended myeloid cell analyses in the BAL of WT mice 8 hours post treatment with cytokine cocktails**. Numbers of **A)** type 1 conventional dendritic cells (cDC1), **B)** CD11b^-^Ly6C^+^ DCs, **C)** CD11b^+^Ly6C^+^ DCs, **D)** CD11b+ DCs, **E)** eosinophils, **F)** Ly6C-monocytes, **G)** lymphocytes in female mice with or without IFN-γ treatment. Unpaired t test. Numbers of **H)** cDC1, **I)** CD11b^-^Ly6C^+^ DCs, **J)** CD11b^+^Ly6C^+^ DCs, **K)** CD11b+ DCs, **L)** eosinophil, **M)** Ly6C-monocyte, **N)** lymphocyte in male mice with or without IFN-γ treatment. Unpaired t test. *p* value: ns≤ 0.1234, *≤ 0.0332, **≤ 0.0021, ***≤ 0.0002, ****≤ 0.0001. All data have n≥3 mice, 2 experiment, mean ± SEM.

**Figure S3: Extended myeloid cell analyses in the BAL of WT and STAT1 KO mice 8 hours post treatment with cytokine cocktails.** Numbers of **A)** type 1 conventional dendritic cells (cDC1), **B)** CD11b^-^Ly6C^+^ DCs, **C)** CD11b^+^Ly6C^+^ DCs, **D)** CD11b+ DCs, **E)** eosinophils, **F)** Ly6C-monocytes, **G)** lymphocytes in both WT and STAT1 KO female mice with IFN-γ treatment. Unpaired t test. Numbers of **H)** cDC1, **I)** CD11b^-^Ly6C^+^ DCs, **J)** CD11b^+^Ly6C^+^ DCs, **K)** CD11b+ DCs, **L)** eosinophils, **M)** Ly6C-monocytes, **N)** lymphocytes in both WT and STAT1 KO male mice with IFN-γ treatment. Unpaired t test. *p* value: ns≤ 0.1234, *≤ 0.0332, **≤ 0.0021, ***≤ 0.0002, ****≤ 0.0001. All data have n≥3 mice, 2 experiment, mean ± SEM.

## REFERENCES

1. Domingues HS, Mues M, Lassmann H, Wekerle H, Krishnamoorthy G. Functional and pathogenic differences of Th1 and Th17 cells in experimental autoimmune encephalomyelitis. PLoS One. 2010;5(11):e15531.

2. Harbour SN, Maynard CL, Zindl CL, Schoeb TR, Weaver CT. Th17 cells give rise to Th1 cells that are required for the pathogenesis of colitis. Proc Natl Acad Sci U S A. 2015;112(22):7061–6.

3. Kryczek I, Bruce AT, Gudjonsson JE, Johnston A, Aphale A, Vatan L, et al. Induction of IL-17+ T cell trafficking and development by IFN-gamma: mechanism and pathological relevance in psoriasis. J Immunol. 2008;181(7):4733–41.

4. Murdock BJ, Shreiner AB, McDonald RA, Osterholzer JJ, White ES, Toews GB, et al. Coevolution of TH1, TH2, and TH17 responses during repeated pulmonary exposure to Aspergillus fumigatus conidia. Infect Immun. 2011;79(1):125–35.

5. Fujii C, Kondo T, Ochi H, Okada Y, Hashi Y, Adachi T, et al. Altered T cell phenotypes associated with clinical relapse of multiple sclerosis patients receiving fingolimod therapy. Sci Rep. 2016;6:35314.

6. Strober W, Fuss IJ. Proinflammatory cytokines in the pathogenesis of inflammatory bowel diseases. Gastroenterology. 2011;140(6):1756–67.

7. Kagami S, Rizzo HL, Lee JJ, Koguchi Y, Blauvelt A. Circulating Th17, Th22, and Th1 cells are increased in psoriasis. J Invest Dermatol. 2010;130(5):1373–83.

8. Ravi VR, Korkmaz FT, De Ana CL, Lu L, Shao FZ, Odom CV, et al. Lung CD4(+) resident memory T cells use airway secretory cells to stimulate and regulate onset of allergic airway neutrophilic disease. Cell Rep. 2025;44(3):115294.

9. Larsen JM, Bonefeld CM, Poulsen SS, Geisler C, Skov L. IL-23 and T(H)17-mediated inflammation in human allergic contact dermatitis. J Allergy Clin Immunol. 2009;123(2):486–92.

10. Bacher P, Hohnstein T, Beerbaum E, Rocker M, Blango MG, Kaufmann S, et al. Human Anti-fungal Th17 Immunity and Pathology Rely on Cross-Reactivity against Candida albicans. Cell. 2019;176(6):1340–55 e15.

11. Khader SA, Bell GK, Pearl JE, Fountain JJ, Rangel-Moreno J, Cilley GE, et al. IL-23 and IL-17 in the establishment of protective pulmonary CD4+ T cell responses after vaccination and during Mycobacterium tuberculosis challenge. Nat Immunol. 2007;8(4):369–77.

12. Scriba TJ, Kalsdorf B, Abrahams DA, Isaacs F, Hofmeister J, Black G, et al. Distinct, specific IL-17- and IL-22-producing CD4+ T cell subsets contribute to the human anti-mycobacterial immune response. J Immunol. 2008;180(3):1962–70.

13. Zielinski CE, Mele F, Aschenbrenner D, Jarrossay D, Ronchi F, Gattorno M, et al. Pathogen-induced human TH17 cells produce IFN-gamma or IL-10 and are regulated by IL-1beta. Nature. 2012;484(7395):514–8.

14. Amezcua Vesely MC, Pallis P, Bielecki P, Low JS, Zhao J, Harman CCD, et al. Effector T(H)17 Cells Give Rise to Long-Lived T(RM) Cells that Are Essential for an Immediate Response against Bacterial Infection. Cell. 2019;178(5):1176–88 e15.

15. Hirota K, Duarte JH, Veldhoen M, Hornsby E, Li Y, Cua DJ, et al. Fate mapping of IL-17-producing T cells in inflammatory responses. Nat Immunol. 2011;12(3):255–63.

16. Smith NM, Wasserman GA, Coleman FT, Hilliard KL, Yamamoto K, Lipsitz E, et al. Regionally compartmentalized resident memory T cells mediate naturally acquired protection against pneumococcal pneumonia. Mucosal Immunol. 2018;11(1):220–35.

17. Yeh WI, McWilliams IL, Harrington LE. IFNgamma inhibits Th17 differentiation and function via Tbet-dependent and Tbet-independent mechanisms. J Neuroimmunol. 2014;267(1-2):20–7.

18. Toh ML, Kawashima M, Zrioual S, Hot A, Miossec P, Miossec P. IL-17 inhibits human Th1 differentiation through IL-12R beta 2 downregulation. Cytokine. 2009;48(3):226–30.

19. Cao H, Diao J, Liu H, Liu S, Liu J, Yuan J, et al. The Pathogenicity and Synergistic Action of Th1 and Th17 Cells in Inflammatory Bowel Diseases. Inflamm Bowel Dis. 2023;29(5):818–29.

20. Komiyama Y, Nakae S, Matsuki T, Nambu A, Ishigame H, Kakuta S, et al. IL-17 plays an important role in the development of experimental autoimmune encephalomyelitis. J Immunol. 2006;177(1):566–73.

21. Leung S, Liu X, Fang L, Chen X, Guo T, Zhang J. The cytokine milieu in the interplay of pathogenic Th1/Th17 cells and regulatory T cells in autoimmune disease. Cell Mol Immunol. 2010;7(3):182–9.

22. Nindl V, Maier R, Ratering D, De Giuli R, Zust R, Thiel V, et al. Cooperation of Th1 and Th17 cells determines transition from autoimmune myocarditis to dilated cardiomyopathy. Eur J Immunol. 2012;42(9):2311–21.

23. Mitsi E, Carniel B, Reiné J, Rylance J, Zaidi S, Soares-Schanoski A, et al. Nasal Pneumococcal Density Is Associated with Microaspiration and Heightened Human Alveolar Macrophage Responsiveness to Bacterial Pathogens. Am J Respir Crit Care Med. 2020;201(3):335–47.

24. Weiser JN, Ferreira DM, Paton JC. Streptococcus pneumoniae: transmission, colonization and invasion. Nat Rev Microbiol. 2018;16(6):355–67.

25. Hill PC, Townend J, Antonio M, Akisanya B, Ebruke C, Lahai G, et al. Transmission of Streptococcus pneumoniae in rural Gambian villages: a longitudinal study. Clin Infect Dis. 2010;50(11):1468–76.

26. Hussain M, Melegaro A, Pebody RG, George R, Edmunds WJ, Talukdar R, et al. A longitudinal household study of Streptococcus pneumoniae nasopharyngeal carriage in a UK setting. Epidemiol Infect. 2005;133(5):891–8.

27. Lipsitch M, Abdullahi O, D’Amour A, Xie W, Weinberger DM, Tchetgen Tchetgen E, et al. Estimating rates of carriage acquisition and clearance and competitive ability for pneumococcal serotypes in Kenya with a Markov transition model. Epidemiology. 2012;23(4):510–9.

28. Regev-Yochay G, Raz M, Dagan R, Porat N, Shainberg B, Pinco E, et al. Nasopharyngeal carriage of Streptococcus pneumoniae by adults and children in community and family settings. Clin Infect Dis. 2004;38(5):632–9.

29. Sibale LL, Lo SW, Kalata N, Nyazika TK, Mitole N, Dyster V, et al. Within-host genetic diversity of pneumococcal serotype 3 during one-year prolonged carriage in a healthy adult. Nat Commun. 2025;16(1):8920.

30. Shenoy AT, Lyon De Ana C, Arafa EI, Salwig I, Barker KA, Korkmaz FT, et al. Antigen presentation by lung epithelial cells directs CD4(+) T(RM) cell function and regulates barrier immunity. Nat Commun. 2021;12(1):5834.

31. Croucher NJ, Kagedan L, Thompson CM, Parkhill J, Bentley SD, Finkelstein JA, et al. Selective and genetic constraints on pneumococcal serotype switching. PLoS Genet. 2015;11(3):e1005095.

32. Sabharwal V, Stevenson A, Figueira M, Orthopoulos G, Trzciński K, Pelton SI. Capsular switching as a strategy to increase pneumococcal virulence in experimental otitis media model. Microbes Infect. 2014;16(4):292–9.

33. Van Bambeke F, Reinert RR, Appelbaum PC, Tulkens PM, Peetermans WE. Multidrug-resistant Streptococcus pneumoniae infections: current and future therapeutic options. Drugs. 2007;67(16):2355–82.

34. Shenoy AT, Wasserman GA, Arafa EI, Wooten AK, Smith NMS, Martin IMC, et al. Lung CD4(+) resident memory T cells remodel epithelial responses to accelerate neutrophil recruitment during pneumonia. Mucosal Immunol. 2020;13(2):334–43.

35. Ahlin A, Elinder G, Palmblad J. Dose-dependent enhancements by interferon-gamma on functional responses of neutrophils from chronic granulomatous disease patients. Blood. 1997;89(9):3396–401.

36. Kirkwood JM, Ernstoff MS, Trautman T, Hebert G, Nishida Y, Davis CA, et al. In vivo biological response to recombinant interferon-gamma during a phase I dose-response trial in patients with metastatic melanoma. J Clin Oncol. 1990;8(6):1070–82.

37. McGinley AM, Sutton CE, Edwards SC, Leane CM, DeCourcey J, Teijeiro A, et al. Interleukin-17A Serves a Priming Role in Autoimmunity by Recruiting IL-1beta-Producing Myeloid Cells that Promote Pathogenic T Cells. Immunity. 2020;52(2):342–56 e6.

38. Sergejeva S, Ivanov S, Lötvall J, Lindén A. Interleukin-17 as a recruitment and survival factor for airway macrophages in allergic airway inflammation. Am J Respir Cell Mol Biol. 2005;33(3):248–53.

39. Shuai K, Horvath CM, Huang LH, Qureshi SA, Cowburn D, Darnell JE, Jr. Interferon activation of the transcription factor Stat91 involves dimerization through SH2-phosphotyrosyl peptide interactions. Cell. 1994;76(5):821–8.

40. Iwanaga N, Chen K, Yang H, Lu S, Hoffmann JP, Wanek A, et al. Vaccine-driven lung TRM cells provide immunity against Klebsiella via fibroblast IL-17R signaling. Sci Immunol. 2021;6(63):eabf1198.

41. Carvalho A, Giovannini G, De Luca A, D’Angelo C, Casagrande A, Iannitti RG, et al. Dectin-1 isoforms contribute to distinct Th1/Th17 cell activation in mucosal candidiasis. Cell Mol Immunol. 2012;9(3):276–86.

42. van de Veerdonk FL, Joosten LA, Shaw PJ, Smeekens SP, Malireddi RK, van der Meer JW, et al. The inflammasome drives protective Th1 and Th17 cellular responses in disseminated candidiasis. Eur J Immunol. 2011;41(8):2260–8.

43. Clemente AM, Castronovo G, Antonelli A, D’Andrea MM, Tanturli M, Perissi E, et al. Differential Th17 response induced by the two clades of the pandemic ST258 Klebsiella pneumoniae clonal lineages producing KPC-type carbapenemase. PLoS One. 2017;12(6):e0178847.

44. Diani M, Altomare G, Reali E. T Helper Cell Subsets in Clinical Manifestations of Psoriasis. J Immunol Res. 2016;2016:7692024.

45. Nistala K, Adams S, Cambrook H, Ursu S, Olivito B, de Jager W, et al. Th17 plasticity in human autoimmune arthritis is driven by the inflammatory environment. Proc Natl Acad Sci U S A. 2010;107(33):14751–6.

46. Schmidt-Weber CB, Akdis M, Akdis CA. TH17 cells in the big picture of immunology. J Allergy Clin Immunol. 2007;120(2):247–54.

47. Lambrecht BN, Hammad H. The immunology of asthma. Nat Immunol. 2015;16(1):45–56.

48. Paun A, Bergeron ME, Haston CK. The Th1/Th17 balance dictates the fibrosis response in murine radiation-induced lung disease. Sci Rep. 2017;7(1):11586.

49. Kamiya M, Carter H, Espindola MS, Doyle TJ, Lee JS, Merriam LT, et al. Immune mechanisms in fibrotic interstitial lung disease. Cell. 2024;187(14):3506–30.

50. Chu CQ, Wittmer S, Dalton DK. Failure to suppress the expansion of the activated CD4 T cell population in interferon gamma-deficient mice leads to exacerbation of experimental autoimmune encephalomyelitis. J Exp Med. 2000;192(1):123–8.

51. Desvignes L, Ernst JD. Interferon-gamma-responsive nonhematopoietic cells regulate the immune response to Mycobacterium tuberculosis. Immunity. 2009;31(6):974–85.

52. Nandi B, Behar SM. Regulation of neutrophils by interferon-γ limits lung inflammation during tuberculosis infection. J Exp Med. 2011;208(11):2251–62.

53. Gonzalez-Ferrer S, Penaloza HF, van der Geest R, Xiong Z, Gheware A, Tabary M, et al. STAT1 Employs Myeloid Cell-Extrinsic Mechanisms to Regulate the Neutrophil Response and Provide Protection against Invasive Klebsiella pneumoniae Lung Infection. Immunohorizons. 2024;8(1):122–35.

54. Duong TT, St Louis J, Gilbert JJ, Finkelman FD, Strejan GH. Effect of anti-interferon-gamma and anti-interleukin-2 monoclonal antibody treatment on the development of actively and passively induced experimental allergic encephalomyelitis in the SJL/J mouse. J Neuroimmunol. 1992;36(2-3):105–15.

55. Eren Akarcan S, Ulusoy Severcan E, Edeer Karaca N, Isik E, Aksu G, Migaud M, et al. Gain-of-Function Mutations in STAT1: A Recently Defined Cause for Chronic Mucocutaneous Candidiasis Disease Mimicking Combined Immunodeficiencies. Case Reports Immunol. 2017;2017:2846928.

56. Okada S, Asano T, Moriya K, Boisson-Dupuis S, Kobayashi M, Casanova JL, et al. Human STAT1 Gain-of-Function Heterozygous Mutations: Chronic Mucocutaneous Candidiasis and Type I Interferonopathy. J Clin Immunol. 2020;40(8):1065–81.

57. Toubiana J, Okada S, Hiller J, Oleastro M, Lagos Gomez M, Aldave Becerra JC, et al. Heterozygous STAT1 gain-of-function mutations underlie an unexpectedly broad clinical phenotype. Blood. 2016;127(25):3154–64.

58. Zhang W, Chen X, Gao G, Xing S, Zhou L, Tang X, et al. Clinical Relevance of Gain- and Loss-of-Function Germline Mutations in STAT1: A Systematic Review. Front Immunol. 2021;12:654406.

59. Moriya K, Suzuki T, Uchida N, Nakano T, Katayama S, Irie M, et al. Ruxolitinib treatment of a patient with steroid-dependent severe autoimmunity due to STAT1 gain-of-function mutation. Int J Hematol. 2020;112(2):258–62.

60. Youakim A, Ahdieh M. Interferon-gamma decreases barrier function in T84 cells by reducing ZO-1 levels and disrupting apical actin. Am J Physiol. 1999;276(5):G1279–88.

61. Bruewer M, Utech M, Ivanov AI, Hopkins AM, Parkos CA, Nusrat A. Interferon-gamma induces internalization of epithelial tight junction proteins via a macropinocytosis-like process. Faseb j. 2005;19(8):923–33.

62. Langer V, Vivi E, Regensburger D, Winkler TH, Waldner MJ, Rath T, et al. IFN-γ drives inflammatory bowel disease pathogenesis through VE-cadherin-directed vascular barrier disruption. J Clin Invest. 2019;129(11):4691–707.

63. Caturegli P, Hejazi M, Suzuki K, Dohan O, Carrasco N, Kohn LD, et al. Hypothyroidism in transgenic mice expressing IFN-gamma in the thyroid. Proc Natl Acad Sci U S A. 2000;97(4):1719–24.

64. Huang Z, Zhang J, Chen S, Mao M, Wang B. Expression levels of interleukin-17 and interferon-γ in peripheral blood and relationship with thyroid function in patients with Hashimoto’s thyroiditis. Front Mol Biosci. 2025;12:1645736.

65. von Herrath MG, Oldstone MB. Interferon-gamma is essential for destruction of beta cells and development of insulin-dependent diabetes mellitus. J Exp Med. 1997;185(3):531–9.

66. Darnell JE, Jr. STATs and gene regulation. Science. 1997;277(5332):1630–5.

67. Broggi A, Granucci F, Zanoni I. Type III interferons: Balancing tissue tolerance and resistance to pathogen invasion. J Exp Med. 2020;217(1).

68. Blazek K, Eames HL, Weiss M, Byrne AJ, Perocheau D, Pease JE, et al. IFN-λ resolves inflammation via suppression of neutrophil infiltration and IL-1β production. J Exp Med. 2015;212(6):845–53.

69. Filipi M, Jack S. Interferons in the Treatment of Multiple Sclerosis: A Clinical Efficacy, Safety, and Tolerability Update. Int J MS Care. 2020;22(4):165–72.

70. Stock AT, Smith JM, Carbone FR. Type I IFN suppresses Cxcr2 driven neutrophil recruitment into the sensory ganglia during viral infection. J Exp Med. 2014;211(5):751–9.

71. Broggi A, Ghosh S, Sposito B, Spreafico R, Balzarini F, Lo Cascio A, et al. Type III interferons disrupt the lung epithelial barrier upon viral recognition. Science. 2020;369(6504):706–12.

72. Major J, Crotta S, Llorian M, McCabe TM, Gad HH, Priestnall SL, et al. Type I and III interferons disrupt lung epithelial repair during recovery from viral infection. Science. 2020;369(6504):712–7.

73. Snell LM, McGaha TL, Brooks DG. Type I Interferon in Chronic Virus Infection and Cancer. Trends Immunol. 2017;38(8):542–57.

74. Chasset F, Dayer JM, Chizzolini C. Type I Interferons in Systemic Autoimmune Diseases: Distinguishing Between Afferent and Efferent Functions for Precision Medicine and Individualized Treatment. Front Pharmacol. 2021;12:633821.

75. Bastard P, Rosen LB, Zhang Q, Michailidis E, Hoffmann HH, Zhang Y, et al. Autoantibodies against type I IFNs in patients with life-threatening COVID-19. Science. 2020;370(6515).

76. Hsieh LL, Thompson EA, Jairam NP, Roznik K, Figueroa A, Aytenfisu T, et al. SARS-CoV-2 induces neutrophil degranulation and differentiation into myeloid-derived suppressor cells associated with severe COVID-19. Sci Transl Med. 2025;17(799):eadn7527.

